# A high-throughput whole cell screen to identify inhibitors of *Mycobacterium tuberculosis*

**DOI:** 10.1101/429423

**Authors:** Juliane Ollinger, Anuradha Kumar, David M. Roberts, Mai A. Bailey, Allen Casey, Tanya Parish

## Abstract

Tuberculosis is a disease of global importance for which novel drugs are urgently required. We developed a whole-cell phenotypic screen which can be used to identify inhibitors of *Mycobacterium tuberculosis* growth. We used recombinant strains of virulent *M. tuberculosis* which express far-red fluorescent reporters and used fluorescence to monitor growth *in vitro*. We optimized our high throughput assays using both 96-well and 384-well plates; both formats gave assays which met stringent reproducibility and robustness tests. We screened a compound set of 1105 chemically diverse compounds previously shown to be active against *M. tuberculosis* and identified primary hits which showed ≥ 90% growth inhibition. We ranked hits and identified three chemical classes of interest – the phenoxyalkylbenzamidazoles, the benzothiophene 1–1 dioxides, and the piperidinamines. These new compound classes may serve as starting points for the development of new series of inhibitors that prevent the growth of *M. tuberculosis*. This assay can be used for further screening, or could easily be adapted to other strains of *M. tuberculosis*.

## Introduction

Tuberculosis (TB), caused by the bacterial pathogen *Mycobacterium tuberculosis*, is a disease of global importance which killed approximately 1.7 million people in 2016 (1). A lengthy 4-drug regimen is used to treat active infection, but drug resistant strains have emerged and threaten efforts to control the disease. Multi-drug resistant (MDR) and extremely drug resistant (XDR) TB are gaining footholds in areas where HIV is predominant and/or antibiotic treatment of patients is administered incompletely or incorrectly (1). Thus, there is an urgent need for new drugs that are effective at killing *M. tuberculosis* and which might shorten therapy.

High throughput screening of small molecules has the potential to identify new compound classes that are effective against *M. tuberculosis*. Biochemical screens have been used to find inhibitors of specific targets, normally essential enzymes. A number of targets have been tested (2–15) but this approach had limited success in finding hits with whole cell activity for a variety of reasons including lack of permeation, efflux, and poor target vulnerability or engagement (16).

In contrast, phenotypic screening relies on identifying compounds with whole cell activity from the outset, with no knowledge of the cellular target. Although there remain challenges in dealing with an organism which grows very slowly and requires handling within specialized containment facilities. A number of assays have been developed which use different approaches, for example the use of non-pathogenic surrogates such as

*Mycobacterium tuberculosis H37Ra* (17), *Mycobacterium smegmatis* (18) or *Mycobacterium aurum* (19). High throughput screening has been conducted with *M. tuberculosis* under a variety of conditions, including nutrient starvation (20), under multiple stresses (21, 22), or during infection of host cells (23, 24). Assays using live cells are also available to determine disruption of specific pathways, such as ATP homeostasis (25), pH homeostasis (26), biofilm formation (27) or under specific conditions such as low oxygen (28).

## Material and Methods

### Bacterial strains and growth conditions

*M. tuberculosis* H37Rv LP (ATCC 25618) (29) was grown in Middlebrook 7H9 medium supplemented with 10% v/v oleic acid, albumin, dextrose, catalase (OADC; Becton Dickinson), 0.05% w/v Tween 80 (7H9-OADC-Tw), and 50 μg/mL hygromycin (7H9-OADC-Tw-hyg), where required. Large scale cultures were grown in 100 mL of medium in 450 cm^2^ roller bottles at 37°C and 100 rpm. *M. tuberculosis* strain CHEAM3 and DREAM8 expressing codon-optimized mCherry and DsRed from plasmids pCherry3 (30) and pBlazeC8 (31), respectively, were used.

### Preparation of assay plates

Medium and compound was dispensed into sterile, black, 384-well, clear bottom plates (Greiner) using a Minitrak (Packard BioScience) with a 384-well head contained in a custom HEPA enclosure. Controls were 100 μM rifampicin in column 1 (final assay concentration of 2 μM rifampicin), DMSO in column 2 (final assay concentration 2%) and 125 nM rifampicin in column 23 (final assay concentration of 2.5 nM). *M. tuberculosis* culture was added to columns 1–23 using a MultiDrop Combi (Thermo Fisher); column 24 was not inoculated (contamination control).

### Growth in plates

*M. tuberculosis* was grown to logarithmic phase (OD_590_ = 0.6–0.9) and filtered through a 0.5 μm cellulose-acetate membrane filter, diluted in fresh medium, and inoculated into 96-well or 384-well plates containing medium. Plates were incubated in plastic bags in a humidified incubator at 37°C. OD and fluorescence were read using a Synergy 4 plate reader (BioTek) with excitation/emission of 586nm/614nm for mCherry and 560nm/590nm for DsRed.

### Data Analysis

OD_590_ and fluorescence readouts were analyzed independently. The coefficient of variation (CV) was calculated as the standard deviation (StdDev) ÷ Mean. For each plate the minimum and maximum growth controls were used to determine the signal to background (S:B) ratio (calculated as MeanMaxSignal ÷ MeanMinSignal), signal to noise (S:N) ratio (calculated as (MeanMaxSignal-MeanMinSignal) ÷ StdDevMinSignal), and the Z’ of the controls (1-((3*StdDevMaxSignal+3*StdDevMinSignal) ÷ (MeanMaxSignal-MeanMinSignal)). For each well, the % inhibition was calculated with reference to the maximum growth control (DMSO only).

The complete data set is available at https://pubchem.ncbi.nlm.nih.gov/bioassay/1259417?viewcode=51D85DCA-C8B4-48D9-B4BE-37687D75149B

## Results and Discussion

### Assay development

We were interested in developing a simple whole cell screen which could be used in multiple formats to assess the anti-tubercular activity of large compound sets. We previously developed an assay to monitor growth based on fluorescence and optical density using a strain of *M. tuberculosis* constitutively expressing the far-red reporter mCherry which was robust and reproducible in 96-well format (32). In this study we used recombinant *M. tuberculosis* constitutively expressing either codon-optimized DsRed or mCherry to develop a 384-well high throughput assay.

We determined the minimum inhibitory concentration (MIC) for rifampicin against the parental strain (H37Rv-LP; ATCC 25618) and both the fluorescent strains (CHEAM3 expressing mCherry from plasmid pCherry3, and DREAM8 expressing DsRed from plasmid pBlazeC8). We used a 10-point, 2-fold serial dilution in independent experiments in 96-well plates. Growth inhibition was calculated compared to control wells (DMSO), and curves fit using the Levenberg-Marqardt algorithm. For both strains, we calculated MICs using OD_590_ and fluorescence independently and observed that the MIC for rifampicin was equivalent to the parental strain H37Rv-LP strain (Table 1). MICs derived using fluorescence as a readout were equivalent to OD_590_-derived values (Table 1). Once we had confirmed the equivalence of the three strains in 96-well plates, we determined key parameters for transferring the assay to higher throughput in 384-well plates.

**Table 1.**
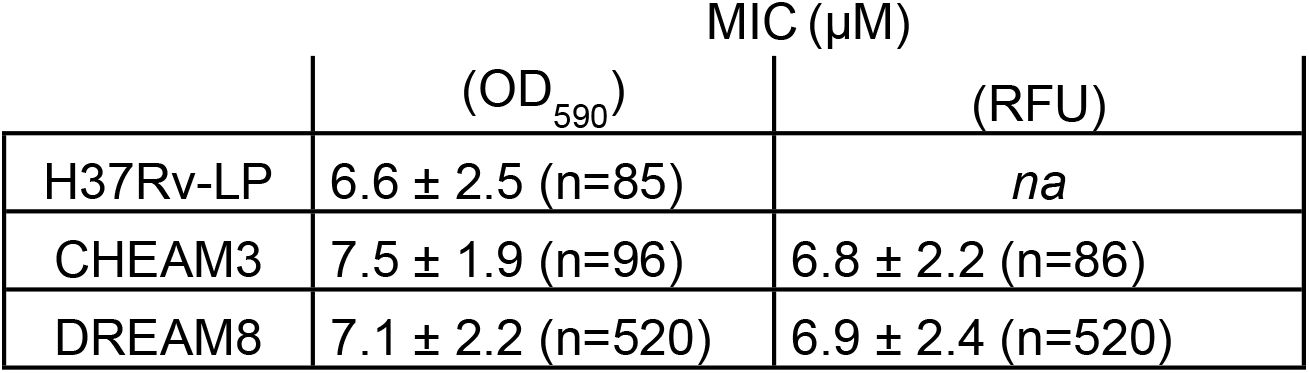
Determination of rifampicin activity against recombinant strains.

| | MIC (μM) | |
| --- | --- | --- |
|  | (OD<sub>590</sub>) | (RFU) |
| H37Rv-LP | 6.6 ± 2.5 (n=85) | *na* |
| CHEAM3 | 7.5 ± 1.9 (n=96) | 6.8 ± 2.2 (n=86) |
| DREAM8 | 7.1 ± 2.2 (n=520) | 6.9 ± 2.4 (n=520) |

MIC, the concentration required to inhibit growth by 90%; *na*, not applicable

### Optimizing fluorescence measurements

We optimized a number of parameters and variables. We had previously determined the optimal parameters for measuring mCherry fluorescence (32). For DsRed, we ran a set of spectral scans varying the excitation wavelength from 540 nm to 565 nm with a fixed emission of 590 nm, and varying the emission wavelength from 580 to 630 nm with a fixed excitation of 558 nm (Figs 1A and B). The signal was optimal at a range of excitation wavelengths around 560 nm, while it peaked at the emission wavelength of 592nm. Based on these scans we selected excitation and emission wavelengths of 560nm and 590nm for DsRed.

**Fig. 1.**
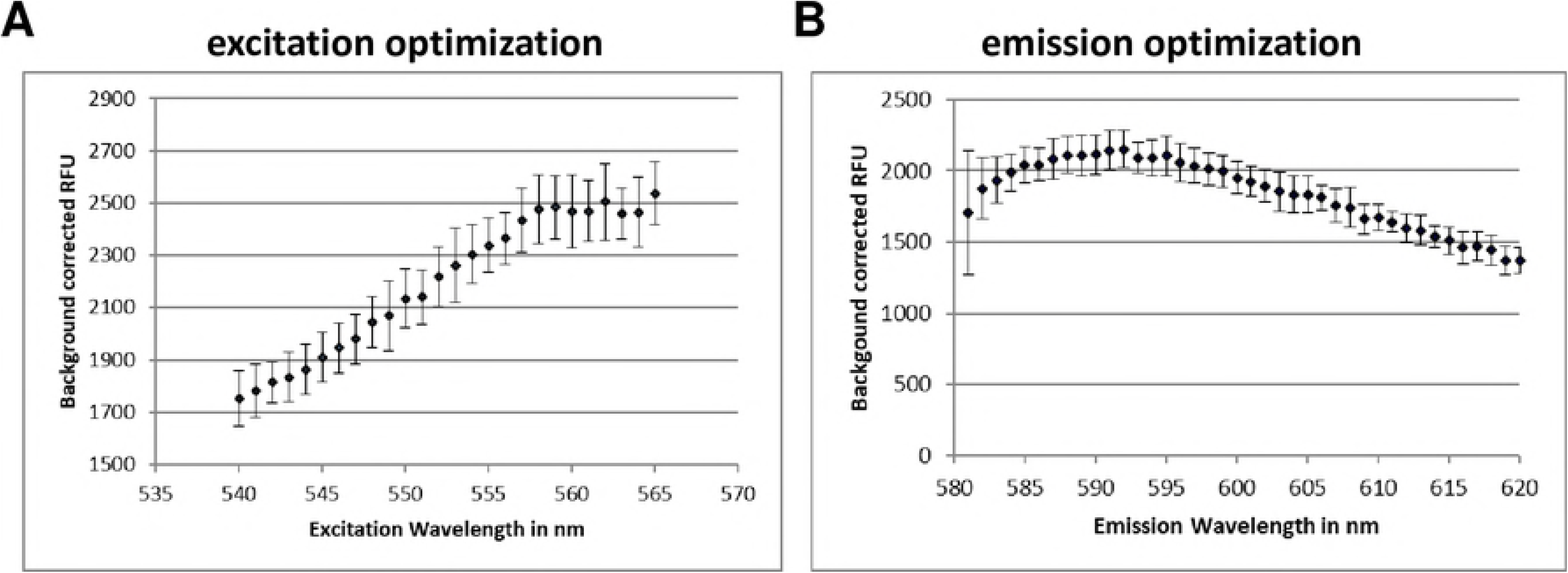
Optimization of fluorescence measurements. DREAM8 was dispensed into 384-well plates and fluorescence was measured at varying excitation wavelengths when the emission wavelength was fixed at 590nm (A) or at varying emission wavelengths when a fixed excitation of 558nm was used (B). Data are the average ± SD from four wells.

We tested the alternatives of bottom and top-read optics in the plate-reader. We compared the use of an external plate shaker or the integral plate shaker in the reader. We obtained the lowest signal to noise (S:N) ratio when plates were not shaken prior to reading and when fluorescence measurements were performed using the bottom optics on the plate reader (data not shown).

### Optimizing inoculum and growth conditions

Several additional assay parameters were tested in the 384-well plates to determine the final assay conditions; the inoculum concentration, and the number of days of incubation prior to measurement of *M. tuberculosis* growth were adjusted to minimize the assay volume while optimizing for signal and reproducibility. At a starting OD_590_ of 0.01, growth was still logarithmic between 4 and 5 days of incubation, whereas at a higher inoculum of OD_590_ = 0.05, cell growth plateaued at 5 days of growth (data not shown). To refine further we performed serial dilutions of CHEAM3 and monitored growth after five days of incubation. We measured fluorescence at the start of the experiment and on day 5 and calculated the S:B ratio (using the values obtained on day 0 of the experiment as the background) (Fig 2A). Wells with starting densities of 0.02 – 0.03 gave the highest S:B (Fig 2A). We ran a similar experiment using starting inocula of 0.01, 0.02 and 0.03 and incubated for 4 or 5 days, but we measured the ratio between full growth and complete inhibition using 2 μM rifampicin (Fig 2B); we found a higher ratio using OD_590_ of 0.02–0.03 (ratio of 13.0 and 13.8 respectively). However, variation was greater using the larger inoculum of OD_590_ = 0.03, with a coeffeicient of variance (CV) of 9%, as compared to a CV of 5% for the inoculum at OD_590_ = 0.02. Thus a starting OD_590_ of 0.02 produced the best signal window and reproducibility in the 384-well assay.

**Fig. 2.**
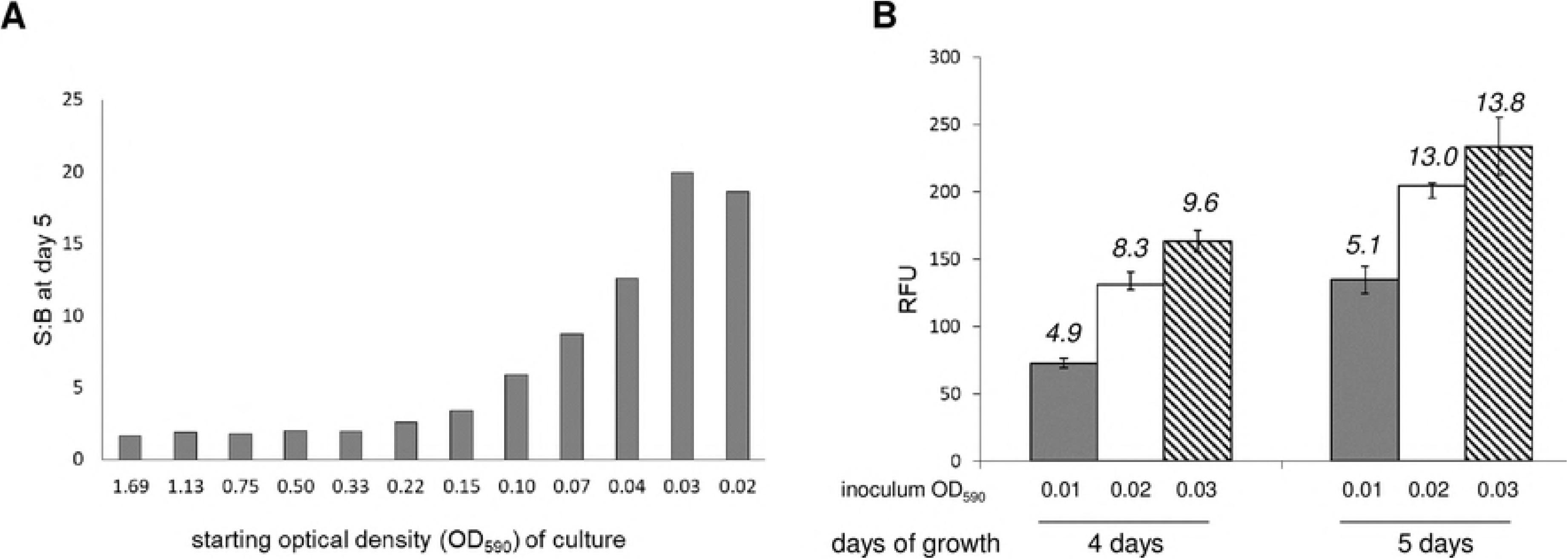
Optimization of growth conditions. (A) Serial dilutions of CHEAM3 were plated in triplicate in a 384 well plate and measured for fluorescence before and after five days of incubation. The signal to background (S:B) for each inoculum was calculated (average signal D0÷average signal D5) to identify the greatest amplitude to measure growth of cells over the course of 5 days. (B) 320 experimental wells of a 384 well plate were inoculated with CHEAM3 diluted to a starting OD_590_ of 0.01, 0.02 or 0.03. Plates were read on D4 and D5. The average fluorescence for each condition (n = 320) was plotted, with error bars indicating the standard deviation. The calculated signal to background (S:B) is shown above each bar.

Our final assay conditions were to inoculate 10 μL of *M. tuberculosis* at OD_590_ = 0.06 into 384-well plates prefilled with 20 μL of medium to give a final theoretical OD_590_ of 0.02. Plates were incubated for 5 days at 37°C and both fluorescence and OD_590_ measured.

### Validation of 384-Well Plate Assay

Once the assay conditions were optimized, we assessed reproducibility according to NCGC guidelines (33). Assay plates containing 20 μL of 7H9-Tw-OADC medium were prepared in a sterile environment. For validation, DMSO or test compounds were added to wells and the plates were inoculated with 10 μL of *M. tuberculosis* at an OD_590_ of 0.06. The final volume in each well was 30 μL, the final OD_590_ was 0.02, and the final concentration of DMSO in each well was 2%. Plates were incubated for five days at 37°C in a humidified incubator. The plate layout was arranged as 320 sample wells in columns 3–22. The remaining four columns were reserved for plate controls: Column 1 – minimum signal (2 μM rifampicin); Column 2 – maximum signal (DMSO); Column 23 – midpoint signal (2.5 nM rifampicin); Column 24 – contamination control (medium only, no inoculum). To test assay reproducibility, we ran a set of six plates independently on three days; two plates of minimum signal, two plates, of maximum signal and two plates of midpoint signal (Fig 3). The % growth in each well was calculated with reference to the maximum signal (Column 2).

**Fig. 3.**
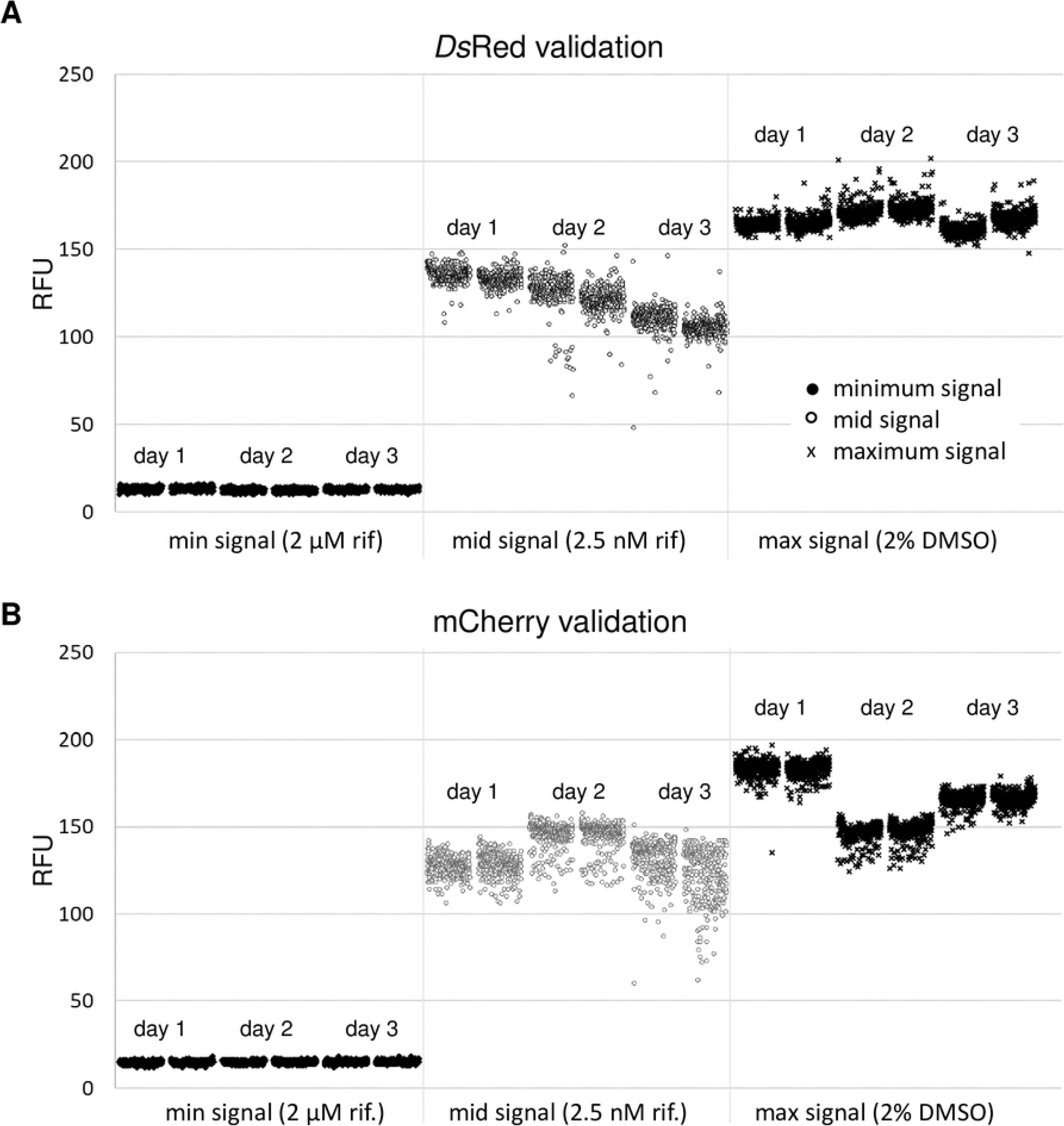
High throughput screen validation scatter plots: Two plates each of containing minimum, midpoint, and maximum signal controls were run in 384-well plates on three separate days using the final assay conditions. Recombinant *M. tuberculosis* expressing (A) DsRed or (B) mCherry was grown for 5 days. Relative fluorescence units (RFU) were measured in each well.

For each strain 2 plates containing maximum signal (Max = *M. tuberculosis* grown with 2% DMSO), mid signal (Mid = *M. tuberculosis* grown in the presence of 2.5 nM rifampicin), and minimum signal (Min = *M. tuberculosis* in the presence of 2 μM rifampicin) were run on three separate days. Max and Min controls (n = 16) from within each plate were used to calculate the signal to noise ratio (S:N), signal to backround ratio (S:B), and Z’ factor (measure of assay robustness as defined earlier). Intraplate controls were also used to calculate the % growth in each well. For the 320 sample wells in each individual plate the average signal, percent coefficient of variance (% CV) of the signal, and average % growth were calculated. The assay statistics generated to validate these two high throughput screens are shown in Tables 2A and B.

**Table 2.** High throughput screen validation statistics. A. DREAM8 validation.

| Signal | Day | Plate | S:N | S:B | Z' | Avg<br>Signal<br>(RFU) | %<br>CV | %<br>growth |
| --- | --- | --- | --- | --- | --- | --- | --- | --- |
| Max | 1 | 1 | 169 | 13 | 0.90 | 12.8 | 6.9 | -0.4 |
| Max | 1 | 2 | 261 | 13 | 0.89 | 12.8 | 6.5 | -0.4 |
| Max | 2 | 1 | 189 | 14 | 0.93 | 13.0 | 8.4 | -0.1 |
| Max | 2 | 2 | 127 | 14 | 0.91 | 13.4 | 7.8 | 0.3 |
| Max | 3 | 1 | 119 | 14 | 0.87 | 12.5 | 7.8 | 0.0 |
| Max | 3 | 2 | 187 | 15 | 0.83 | 12.3 | 7.4 | 0.1 |
| Mid | 1 | 1 | 179 | 13 | 0.89 | 110.6 | 6.4 | 60.7 |
| Mid | 1 | 2 | 250 | 13 | 0.88 | 105.1 | 4.7 | 58.3 |
| Mid | 2 | 1 | 244 | 12 | 0.93 | 135.8 | 3.3 | 79.2 |
| Mid | 2 | 2 | 106 | 12 | 0.92 | 132.8 | 3.2 | 78.4 |
| Mid | 3 | 1 | 226 | 14 | 0.90 | 126.1 | 7.8 | 69.8 |
| Mid | 3 | 2 | 187 | 14 | 0.87 | 121.4 | 5.2 | 68.0 |
| Min | 1 | 1 | 178 | 13 | 0.88 | 161.2 | 2.1 | 97.3 |
| Min | 1 | 2 | 172 | 13 | 0.86 | 167.9 | 2.7 | 96.6 |
| Min | 2 | 1 | 136 | 12 | 0.87 | 164.8 | 1.7 | 97.0 |
| Min | 2 | 2 | 167 | 12 | 0.94 | 165.4 | 2.3 | 97.6 |
| Min | 3 | 1 | 180 | 13 | 0.86 | 171.5 | 2.8 | 99.2 |
| Min | 3 | 2 | 149 | 14 | 0.85 | 173.3 | 2.7 | 97.0 |

**B. CHEAM3 validation** *intra-plate controls*
| Signal | Day | Plate | S:N | S:B | Z' | Avg<br>Signal<br>(RFU) | %<br>CV | %<br>growth |
| --- | --- | --- | --- | --- | --- | --- | --- | --- |
| Max | 1 | 1 | 169 | 12 | 0.89 | 14.6 | 6.5 | 0.0 |
| Max | 1 | 2 | 161 | 12 | 0.87 | 14.8 | 6.5 | 0.1 |
| Max | 2 | 1 | 200 | 11 | 0.79 | 14.8 | 6.0 | -0.3 |
| Max | 2 | 2 | 186 | 11 | 0.70 | 15.0 | 6.1 | -0.4 |
| Max | 3 | 1 | 273 | 11 | 0.90 | 15.1 | 6.2 | -0.1 |
| Max | 3 | 2 | 175 | 11 | 0.88 | 15.1 | 6.4 | -0.2 |
| Mid | 1 | 1 | 245 | 12 | 0.88 | 127.2 | 4.5 | 63.4 |
| Mid | 1 | 2 | 172 | 12 | 0.79 | 128.2 | 5.3 | 63.9 |
| Mid | 2 | 1 | 167 | 10 | 0.80 | 145.1 | 5.0 | 90.7 |
| Mid | 2 | 2 | 177 | 10 | 0.75 | 144.9 | 5.6 | 90.5 |
| Mid | 3 | 1 | 150 | 11 | 0.91 | 131.1 | 7.7 | 77.3 |
| Mid | 3 | 2 | 247 | 10 | 0.90 | 122.5 | 12.3 | 68.1 |
| Min | 1 | 1 | 237 | 12 | 0.93 | 183.3 | 2.8 | 100.4 |
| Min | 1 | 2 | 206 | 12 | 0.86 | 183.1 | 2.5 | 99.7 |
| Min | 2 | 1 | 130 | 10 | 0.78 | 145.9 | 6.1 | 98.3 |
| Min | 2 | 2 | 137 | 10 | 0.75 | 147.6 | 3.7 | 101.2 |
| Min | 3 | 1 | 187 | 11 | 0.88 | 165.7 | 2.7 | 98.2 |
| Min | 3 | 2 | 249 | 11 | 0.88 | 166.6 | 2.5 | 97.6 |

The Z’ factor, an indication of the robustness of the assay (34) was ≥ 0.7 in all plates. S:N was > 100 for both strains and S:B was ≥ 12 for DsRed and ≥ 10 for mCherry. The CV was < 20% in all plate. The average mid-point signal did not vary more than 1.5-fold within a run or across the three runs. Thus, both assays passed statistical validation.

### High throughput screen

To examine the performance of our high-throughput assay, we tested a set of compounds with known activity against *M. tuberculosis*. The Tuberculosis Antimicrobial Acquisition and Coordinating Facility (TAACF) at the Southern Research Institute (SRI) screened libraries containing over 300,000 compounds to identify inhibitors of *M. tuberculosis* growth (35, 36). From the hits identified in these two screens a diversity set of 1105 compounds was made obtained from the Division of Microbiology and Infectious Disease (DMID), National Institute of Allergy and Infectious Diseases (NIAID.as a library of potential anti-tubercular agents for the further development of *M. tuberculosis* drug development assays (37). We obtained this set of compounds and tested them in both HTS-validated assays.

Compounds were obtained in plates, diluted to 0.35 mg/mL and transferred directly into assay plates to yield a final assay concentration of 7 μg/mL (final concentration of 2% DMSO). The standard assay conditions were used for each strain and % growth inhibition was plotted (Fig 4A and B). The Pearson coefficient, a statistical measurement of correlation between two data sets, was calculated for each strain. Fig 4A shows the replicate runs of CHEAM3 with a Pearson coefficient of r= 0.9843. When DREAM8 was the strain used in the screen the correlation coefficient was r= 0.9855 (Fig 4B). Thus the assay performed well in repeat runs with a large compound set.

**Fig. 4.**
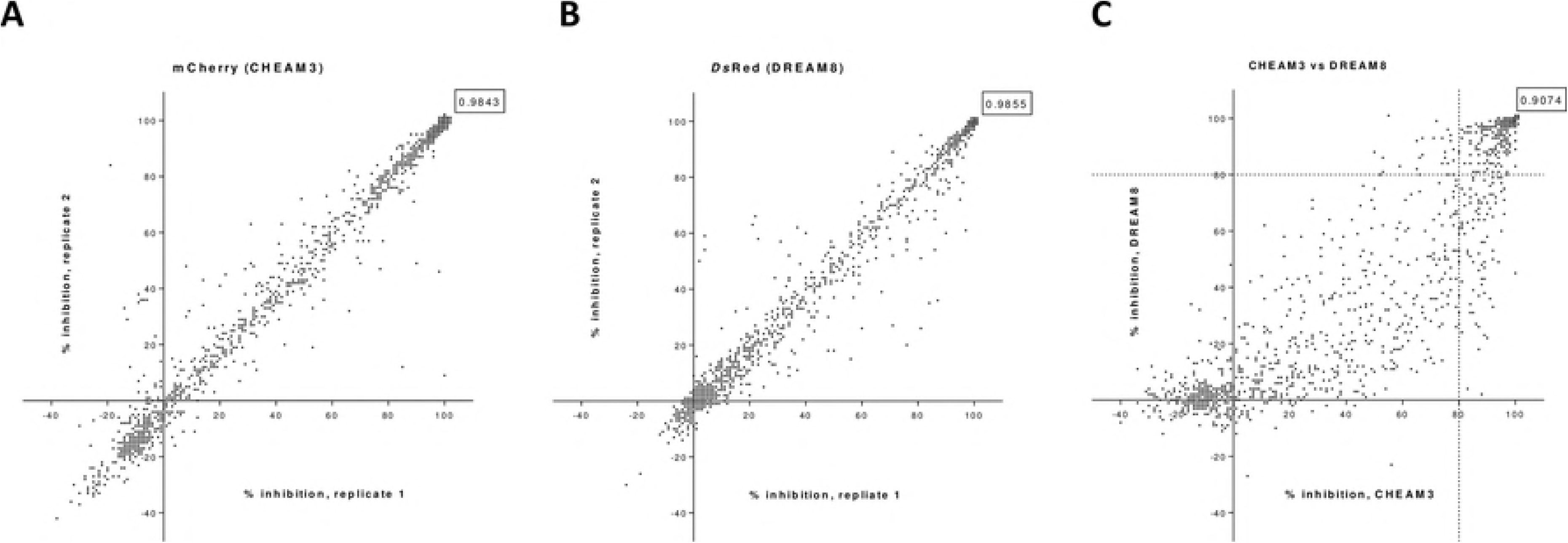
Small molecule compound library screen: A selected library of 1105 small molecules from NIH-SRI/TAACF was screened in replicate experiments against **(A)** CHEAM3 or **(B)** DREAM8. The % inhibition for each compound was calculated. The results from the first and second runs are plotted on the x- and y-axis, respectively, for both strains. For each strain the Pearson coefficient of linear correlation between the two replicate data sets was calculated in Graphpad Prism and is shown in boxed text in the upper right corner of each plot. **(C)** Compounds’ average % inhibition of CHEAM3 and DREAM8 growth was calculated and plotted on the x- and y-axis, respectively. The calculated Pearson co-efficient comparing the data generated from the two different strains is shown in boxed text in the upper right corner of the plot.

We compared the data between the two strains of *M. tuberculosis* strains (Fig 4C). There was a linear relationship between the two strains with a Pearson coefficient of r= 0.9074. Thus, there was no statistical difference between the two strains.

Using CHEAM3, 470 compounds inhibited *M. tuberculosis* growth ≥ 80% while 169 compounds inhibited ≥ 99% (Fig 4A). Using DREAM8, 403 compounds inhibited ≥ 80% growth and 182 compounds inhibited ≥ 99% growth (Fig 4B). 377 compounds inhibited ≥ 80% growth in both strains. There were some minor differences between the two strains. 141 compounds showed ≥ 99% inhibition in both strains, with an additional 69 compounds with %I ≥ 99 in only one of the two strains (41 in DREAM8 and 28 in CHEAM3). However, of these 69 compounds 64 inhibited growth of the alternate strain by at least 90%. The hits from the screen were analysed and revealed three chemotypes shown in Fig 5 that looked most interesting for further development: the phenoxyalkylbenzimidazoles (PAB), the benzothiophene 1–1 dioxides (BTD), and the piperidinamines (PIP).

**Fig. 5.**
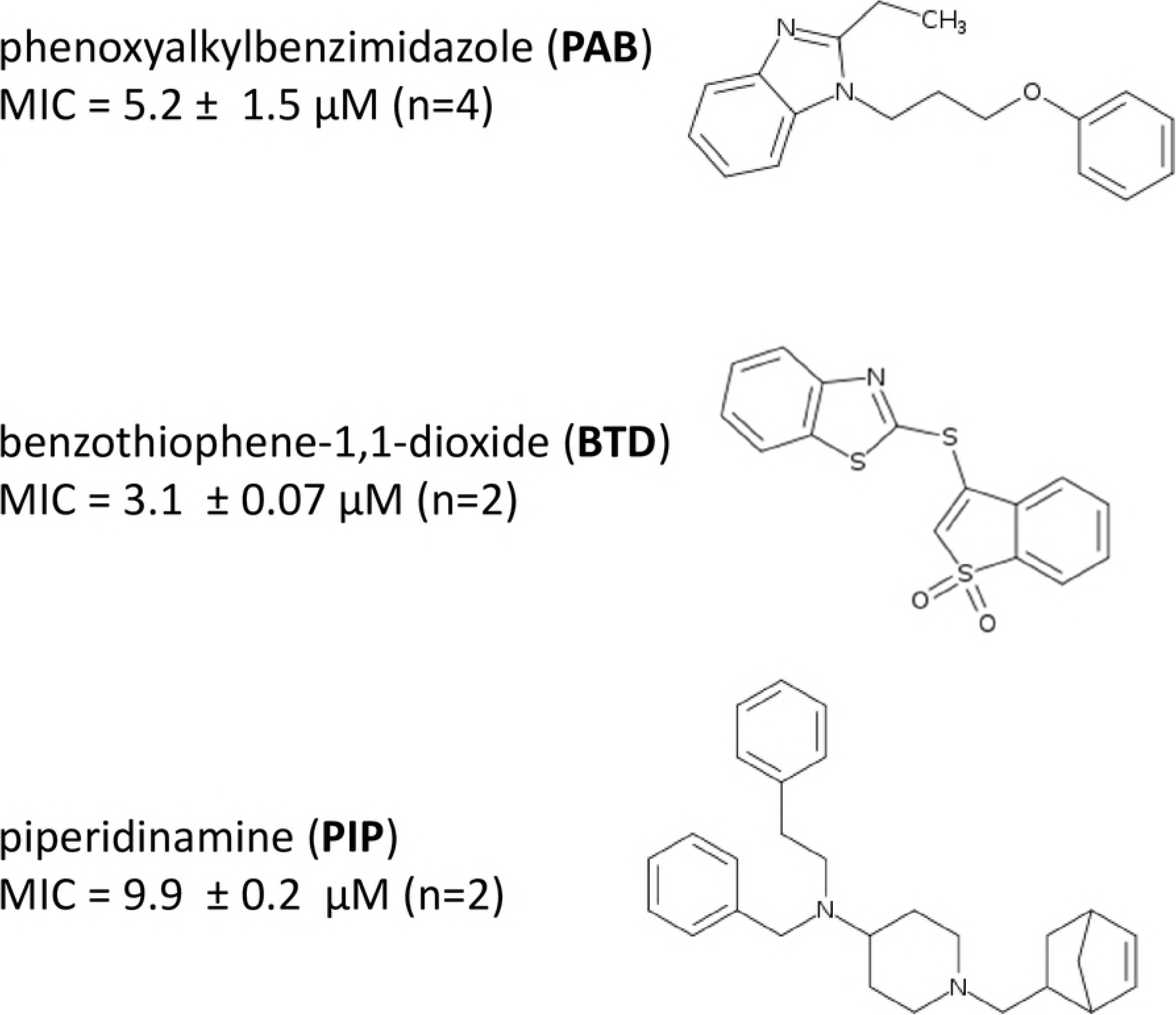
Selected hit compounds from screen: Three hit chemotypes identified in our screen were noted as being especially interesting for further development. Their structures and MICs are shown.

## Discussion

We developed high throughput assays capable of screening large numbers of compounds using two fluorescent reporter strains of *M. tuberculosis*. We used both assays to screen a set of known compounds. Results from these assays were reproducible and the two strains yielded comparable results.

Only a fraction of the compounds we tested had activity against *M. tuberculosis* in our assays, even though these had previously been identified as active (35, 36). A significant difference between the two screens is that we used a lower concentration of 7 μg/mL, as compared to 20 μg/ml previously used; thus we will only detect the more potent compounds. There are also some technical differences between the two assays, in particular that we used OD_590_ and fluorescence as a measure of increase in bacterial numbers, whereas the previous screen used Alamar Blue which monitors metabolic activity.

Using our fluorescent reporter strains, we identified 210 compounds that inhibited ≥ 99% growth, 141 of which inhibited ≥ 99% growth in both strains. We highlighted three chemotypes that were most interesting as the phenoxyalkylbenzimidazoles (PAB), benzothiophene 1–1 dioxides (BTD), and piperidinamines (PIP). We selected these chemical classes as being novel anti-mycobacterials and we (and others) have investigated these further in other publications pertaining to their recent characterization and development.

Compounds containing the benzimidazole core have long been known to have broad spectrum antibacterial activity (38, 39). In the *M. tuberculosis* phenotypic screen performed by Ananthan et al., 88 compounds with the phenoxyalkylimidazole core were tested, PAB being the most potent active identified in this series, and the only compound in that study with benzimidazole substituted for the imidazole (35). There have since been reports of benzimidazoles having activity against *M. tuberculosis* (40, 41). We investigated this series and confirmed its potent activity and selectivity (42). We have also shown that this class of compounds targets the electron transport chain, specifically targeting QcrB (43).

The benzothiophene 1–1 dioxides (BTD) series was also highlighted by Ananthan et al., as a chemotype. There were only a small number of analogs within this chemotype in their larger screen, all containing a thioether group at the 3-position and the most potent being the compound shown in Fig 5. They further noted that there were 40 additional compounds with the thiopene 1–1 dioxides lacking the benzene substitution in their primary screen, all of which lacked activity. Using this as a starting point we explored the BTD series and evaluated their activity against *M. tuberculosis*. We were able to derive compounds that had good anti-tubercular activity (with MICs of 3–8 μM) but were unable to identify potent compounds analogs that were not cytotoxic to eukaryotic cells (44).

Some work has been done on piperidinamine-containing molecules as motilin-receptor agonists (45) and their clinical development for treating type 1 diabetes (46, 47). Our screen identified the piperidinamines (PIP) as a potent chemotype. We obtained analogs within this chemotype that were commercially available and evaluated them for activity against *M. tuberculosis*, but none showed activity (48). Based on our exploration of modifications around the piperidine core, we were unable to identify avenues for improved activity, and to our knowledge, the PIP chemotype has not yet further developed as an anti-tubercular agent.

In summary, we have developed and validated a robust whole cell phenotypic assay for *M. tuberculosis* in 384-well plates using one of two fluorescent reporters with equivalent outcomes. We used these assays to identify potent inhibitors of *M. tuberculosis* growth that have since proven to be interesting leads for drug development. This assay provides robust and reproducible results, and can be a core tool for high-throughput screening of large chemical libraries for the discovery of novel chemical entities to treat tuberculosis

## Acknowledgements

None

